# Metabolic Cost of Rapid Adaptation of Single Yeast Cells

**DOI:** 10.1101/564419

**Authors:** Gabrielle Woronoff, Philippe Nghe, Jean Baudry, Laurent Boitard, Erez Braun, Andrew D. Griffiths, Jérôme Bibette

## Abstract

Cells can rapidly adapt to changing environments through non-genetic processes; however, the metabolic cost of such adaptation has never been considered. Here we demonstrate metabolic coupling in a remarkable, rapid adaptation process (10^-3^cells/hour) by simultaneously measuring metabolism and division of thousands of individual *Saccharomyces cerevisiae* cells using a droplet microfluidic system. Following a severe challenge, most cells, while not dividing, continue to metabolize, displaying a remarkably wide diversity of metabolic trajectories from which adaptation events can be anticipated. Adaptation requires the consumption of a characteristic amount of energy, indicating that it is an active process. The demonstration that metabolic trajectories predict *a priori* adaptation events provides the first evidence of tight energetic coupling between metabolism and regulatory reorganization in adaptation.

**One Sentence Summary:** Demonstration of the tight coupling between metabolic activity and regulatory processes during rapid adaptation at the single-cell level.

## Main Text

How populations of cells adapt to changing environments remains a major question in evolutionary biology. In the classical neo-Darwinian picture, random genetic mutations cause phenotypic variations, enabling adaptation to the new environment. However, adaptation may also be mediated by non-genetic processes (*1*). Phenotypic plasticity can, for example, allow cells to survive severe environmental challenges, which is essential for adaptive evolution (*2*). It is also suggested to play an important role in diverse biological processes, including cell differentiation and the emergence and progression of diseases such as cancer (*3*). However, the connection between metabolic activity and adaptation has remained elusive.

To investigate adaptation of the yeast *S. cerevisiae* confronted with a severe environmental challenge, we genetically “rewired” cells by detaching the essential *HIS3* gene of the histidine biosynthesis system from its native regulatory system and placed it under the control of the *GAL* system, which is highly induced in the presence of galactose and strongly repressed in glucose (*4*). Switching cells from galactose to glucose in a medium lacking histidine presents the yeast with the challenge of reinitiating histidine synthesis in order to resume growth and prevent extinction. The population dynamics of this system have previously been studied in detail (*4–10).* After switching to glucose, growth continues for ~1 day (Phase I) then, after a few days with little or no growth (Phase II), normal growth is resumed at the population level (Phase III) with a doubling time similar to that before the challenge, indicating that after only a few days the population is fully adapted (Fig. 1).

**Fig. 1.**
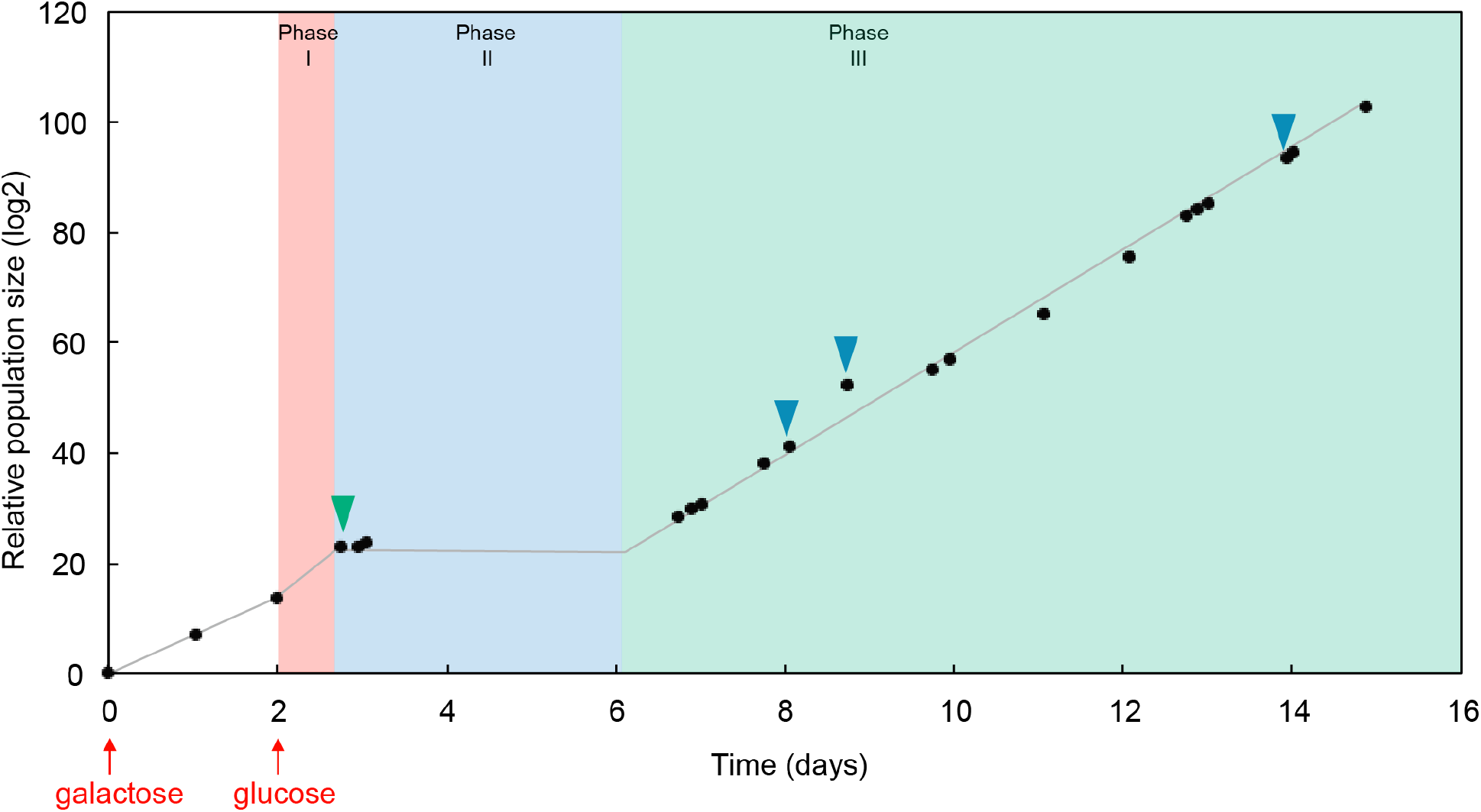
Population growth in bulk cultures during adaptation. The culture of rewired cells started with galactose as the sole carbon source (first red arrow). Then, the medium was switched to glucose (second red arrow). The culture continued to grow during the “initial growth phase” (phase I), before entering the “latent phase” (phase II) where growth stopped. Finally, the population grew again, entering the “adapted phase” (phase III). Green and blue triangles indicate time points when samples were taken for single-cell measurements in droplets (green triangle: 3 hours after the beginning of phase II; blue triangles: 2, 3 and 6 days after the onset of phase III).

To track adaptation at the single-cell level, we used a droplet microfluidic system that allows simultaneous measurements of growth and metabolism of several thousands of single yeast cells over time (*11*). Individual cells, harvested from batch cultures 3 h after the beginning of Phase II (Fig. 1; green triangle), were compartmentalized in 30 pL aqueous droplets in an inert carrier oil that were immobilized in a 2D array in a closed glass observation chamber and incubated at 30°C for 65 to 70 h (Fig. 2A). A small fraction of the droplets contained single cells (typically 6%), and these were surrounded by empty droplets. The consumption of nutrients (glucose) in a cell-containing droplet creates an osmotic imbalance, resulting in water flux, inducing the shrinkage of the droplet and the swelling of neighboring cell-free droplets (Movie S1). Images were taken every 20 or 30 minutes over 3 days and used to determine the change in volume of each droplet as a function of time, reflecting cell metabolism (*11*), while simultaneously imaging the cells to measure division (Materials and Methods). Three independent batch cultures were analyzed, however, only the consolidated data from these three independent experiments is plotted and analyzed (see Table S1 for analysis of individual experiments).

**Fig. 2.**
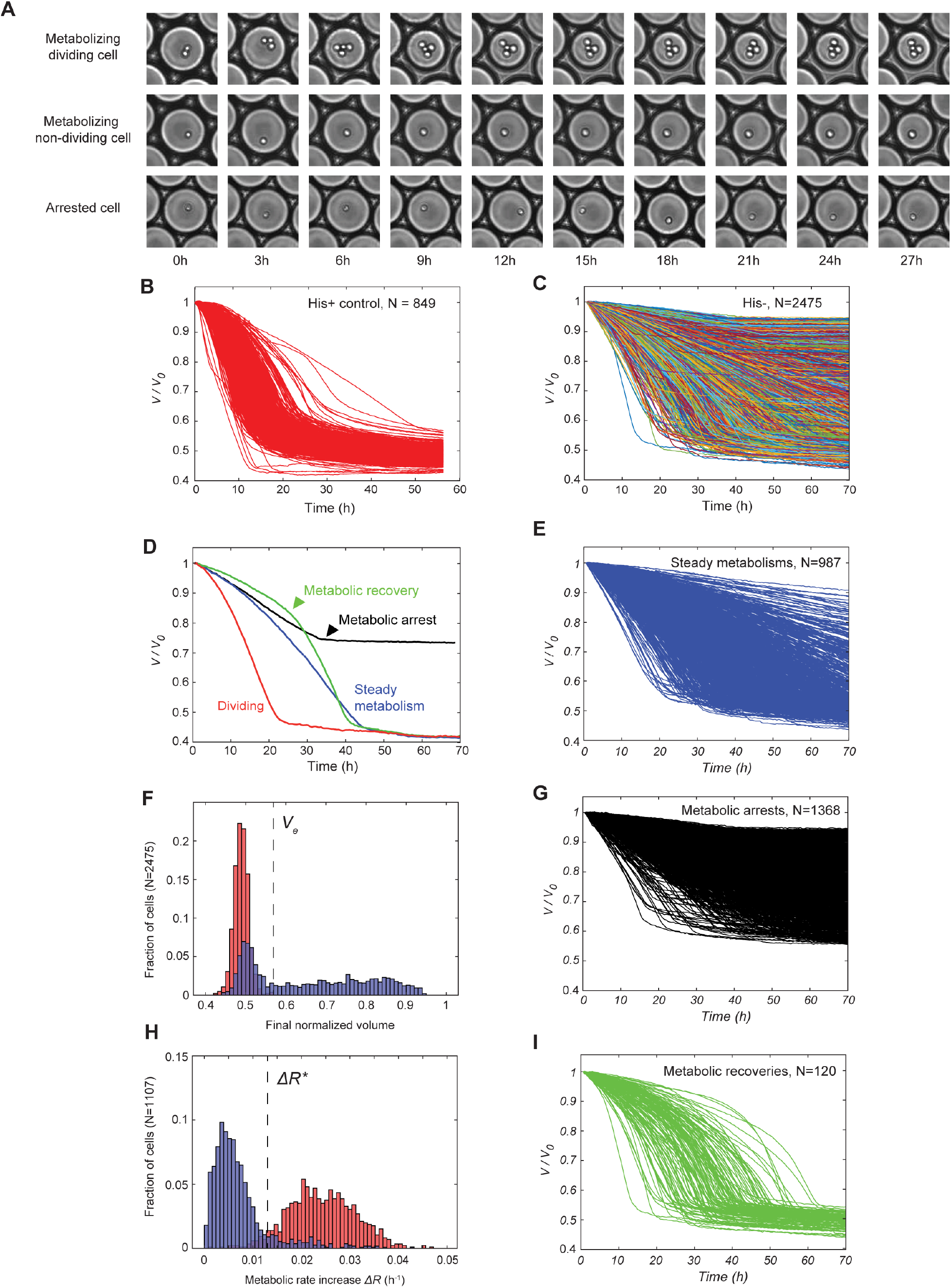
Single-cell analysis of metabolic dynamics in droplets. A) Time-lapse sequences of droplets containing a metabolizing dividing cell, a metabolizing non-dividing cell and an arrested cell. B) Traces of droplet volumes *V* normalized by the initial volume *V_0_*, in the absence of metabolic challenge (with histidine). C) Traces of normalized droplet volumes starting 3 hours after switching from galactose to glucose. Lines are randomly color coded. D) Examples of traces of normalized droplet volumes of each category of metabolic response. E) Subset of steady metabolisms from panel C. F) Distribution of final normalized droplet volume (*V_f_/V_0_*) in the presence (red) and absence (blue) of histidine. *V_e_* indicates the largest final droplet volume with histidine. G) Subset of arrested metabolisms from panel C. H) Distributions of the difference *ΔR* (metabolic rate increase) between the initial (at 2 hours) and maximum metabolic rates in the presence (red) and absence (blue) of histidine. *ΔR*\* indicates the cross-over between the two distributions. I) Subset of metabolic recoveries from panel C.

Metabolic activity was detectable for 88% of cells (final droplet volume <0.95 of initial volume, see Materials and Methods, Fig. S1). The volume variation of droplets containing cells displaying metabolic activity was plotted for both the control experiment, with histidine in the medium (Fig. 2B), and the adaptation experiment, in the absence of histidine (Fig. 2C). Cells in medium lacking histidine showed remarkably diverse metabolic trajectories (CV = 0.60 at 10 hours), while in the presence of histidine the diversity of the metabolic trajectories was more restricted (CV = 0.24 at 10 hours, Fig. S2) and similar to wild-type yeast (*11*).

Analysis of the change of droplet volumes over time (Fig. S1–5) revealed 3 different classes of metabolizing cells (Fig. 2D, Materials and Methods): (i) steadily metabolizing cells (35% of the total population) (Fig. 2E); (ii) cells that underwent metabolic arrest (49% of the total population), the volume curve flattening before nutrient exhaustion, i.e. at a value larger than the final volume of cells grown with histidine (Fig. 2F,G), and (iii) cells that underwent metabolic recovery (4% of the total population), the shrinkage rate increasing to a similar level as for cells under non-stressed conditions (in the presence of histidine) (Fig. 2H,I). The time of metabolic arrest (*T*_arr_) and time of metabolic recovery (*T*_rec_) were determined as the minimum and the maximum of the second derivative of the drop-shrinkage curves, respectively (arrowheads in Fig. 2D, Materials and Methods).

We next addressed the question of the correlation between metabolic profiles and adaptation — the resumption of division. Each metabolizing cell within the shrinking drops was visualized to determine if division occurred at least once and at what time; the time of division *T*_div_ being defined as the time of appearance of a first bud eventually leading to a division event.

We found that within the subset of cells showing steady metabolism, only 9% of the cells resumed division, whereas within the metabolic recovery class, 73% resumed division (Fig. 3A,B). Plotting the distribution of the time difference between the onset of cell division (*T*_div_) and metabolic recovery (*T*_rec_) shows that the two events are strongly correlated (R^2^=0.79, p<10^-5^, Fig. S6), with a mean value *T*_div_ – *T*_rec_ ≈ −1.35 ± 5.5 h (Fig. 3C), which is significantly smaller than the average time of recovery (29 h). This indicates that division starts when the maximum metabolic acceleration is reached.

**Fig. 3.**
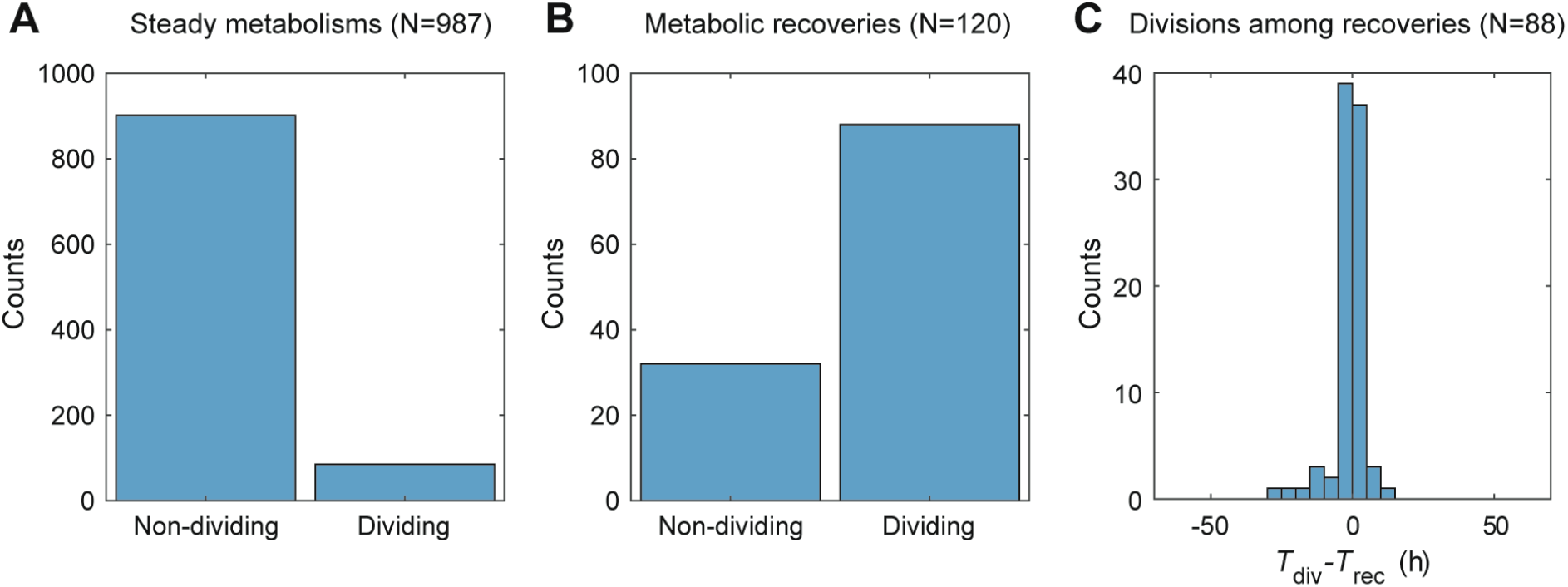
Relation between onset of division and metabolic recovery. A) Fraction of dividing cells in steady metabolisms. B) Fraction of dividing cells in metabolic recoveries. C) Distribution of the difference between division time (*T_div_*) and metabolic recovery time (*T_rec_*).

The cumulated fraction of metabolic recoveries (Fig. 4A, green) and cells re-commencing division (Fig. 4A, red) both displayed a sigmoidal shape, with a quadratic increase up to 30 hours, followed by a leveling off. The instantaneous rate of adaptation also increased up to 30 h, reaching a maximum of ~10^-3^ cells/h (Fig. 4B). Plotting the recovery time (*T*_rec_) as a function of the inverse initial metabolic rate (R_0_^-1^) revealed a clear linear correlation (R^2^=0.33, p<10^-3^, Fig. 4C). The droplet volume on recovery peaks at ~74% of the initial droplet volume (Fig. 4D, Table S1), indicating that, on average, a characteristic amount of glucose (156 pg) is required for a cell to adapt. The distribution of adaptation times is explained by the distribution of the initial metabolic rates which peaks at low values (Fig. 4E, Materials and Methods). In contrast, the cumulated fraction of metabolic arrests increases quadratically throughout the course of the experiments (Fig. 4F), leading to an instantaneous death rate that continuously increases with time (Fig. 4G). The time of metabolic arrest (*T*_arr_) is, however, poorly correlated with the initial metabolic rate (R^2^=0.12, p<10^-3^, Fig. 4H), in agreement with an age driven mechanism essentially independent of the initial metabolic rate.

**Fig. 4.**
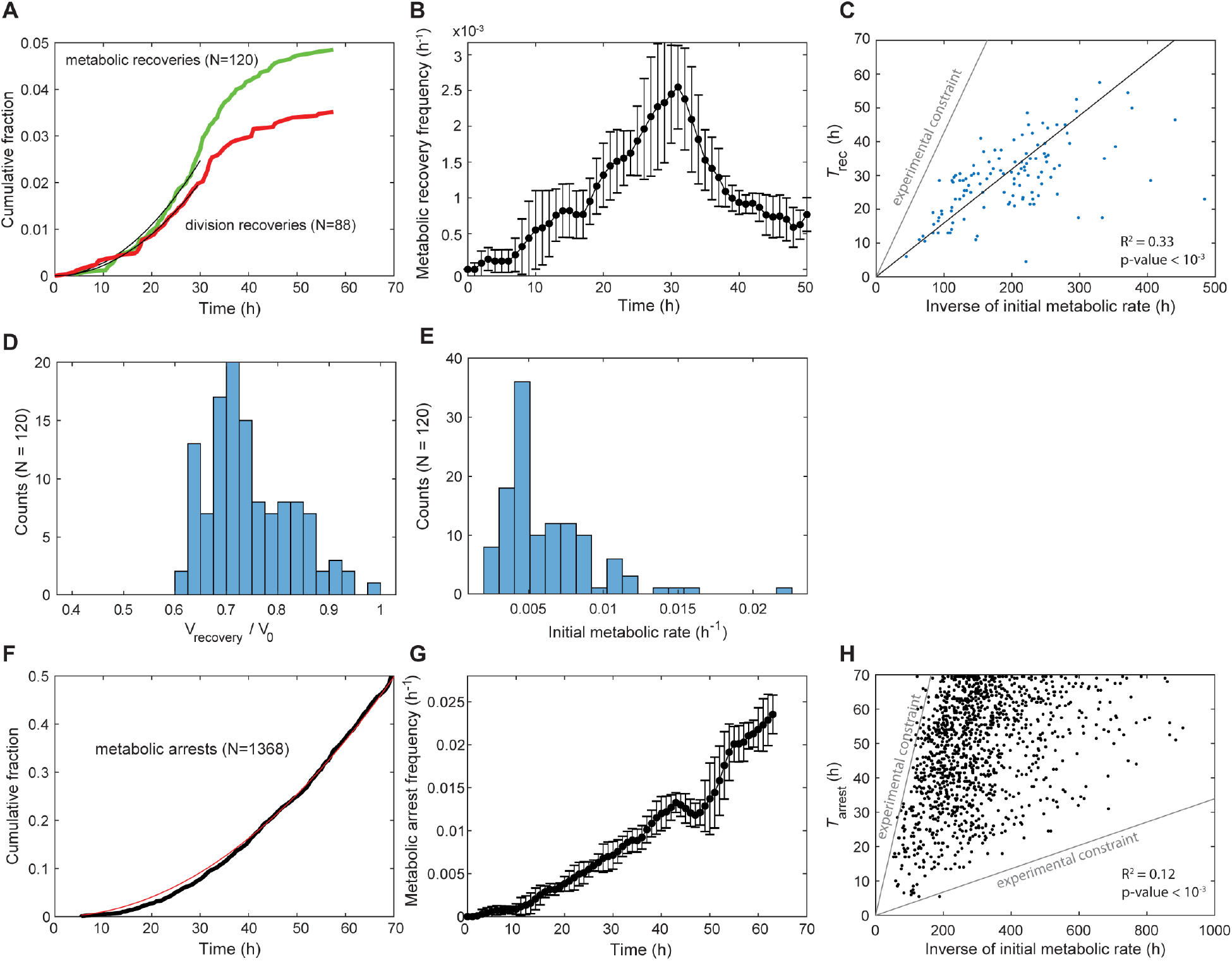
Characterization of adaptation. A) Cumulative fraction over time for metabolic recoveries (green) and division recoveries (red). Black lines are a quadratic fit over the first 30 hours. B) Instantaneous frequency of metabolic recoveries of cells. At each time point, frequencies are computed over 7 consecutive points and error bars are standard deviations over the values obtained over these points. C) Plot of metabolic recovery time (*T_rec_*) as a function of inverse initial metabolic rate (*R*_0_^-1^). The parameter region above the gray line corresponds to cells that have exhausted droplet resources before being able to recover (Material and Methods). D) Distribution of normalized droplet volume at *T_rec_*. E) Distribution of *R*_0_ for the subset of metabolic recoveries. F) Cumulative fraction over time for metabolic arrests (black). The red line is a quadratic fit. G) Instantaneous frequency of arrests of cells that have neither recovered nor arrested their metabolism. Frequencies and error bars were computed as in B). H) Plot of time of metabolic arrest (*T_arr_*) as a function of *R*_0_^-1^. The parameter region above the upper gray line corresponds to cells that have exhausted droplet resources before arresting. The parameter region below the lower gray line corresponds to curves whose final volume is indistinguishable from curves of empty droplets (overall volume variation <5%). The contribution of the constraints in panels C) and H) has been accounted for to compute the coefficients of determination R^2^ (Material and Methods).

We finally examined the metabolism of single cells in the adapted populations taken 2, 4 and 7 days after the batch population entered Phase III (Fig. 1; blue arrows). The overall dynamics of the adapted population (Fig. S7A), is highly comparable with cells in the absence of stress (Fig. 2B), with a similar spread of final droplet volumes (Fig. S7B) and metabolic rates (Fig. S7C, CV=0.35 for adapted compared to CV=0.24 for non-stressed cells). None of the 174 dividing cells analyzed underwent metabolic arrest, demonstrating stability of the adapted state over days, consistent with results at the population level (*4, 9*).

What mechanism might explain this rapid adaptation? Consistent with previous results (*7*), the adaptation process observed here appears to differ from known *S. cerevisiae* stress responses, including the response to amino-acid starvation (12), in a number of ways. Known stress responses are rapid and transient (with changes in gene expression typically peaking after <2 hours) (*13, 14*), and all cells respond broadly similarly (14). In contrast, the characteristic timescale of adaptation of single cells that we observe is much longer (~10^3^ h, see Fig. 4B) and only a fraction of cells adapt. However, the rate of adaptation (~10^-3^ h^-1^) is also several orders of magnitude higher than rates reported based on genetic mechanisms (*15–17*), for example, the spontaneous reversion rates by point mutation in quiescent histidine auxotrophic yeast cells in the absence of histidine is only 10^-5^ to 10^-9^ h^-1^ (*16*). This may point to an epigenetic mechanism, as recently suggested for a similar engineered yeast system (*18*).

In conclusion, this work establishes that, in yeast, rapid adaptation necessitates consumption of a characteristic amount of energy, causing certain cells that metabolize more efficiently to adapt more rapidly. The adapted state is stable, backing up previous observations in bulk, which indicated that the adapted state is stably inherited across generations (*4, 9*), showing that this is a genuine adaptation process. It is also an active process, requiring the consumption of energy, which implies exploration of different states, and fixation of the solution(s) that allow adaptation.

Finally, it is tempting to speculate that a related system may also play a role in other important processes, including cellular differentiation and development. Indeed, stochastic fluctuations at the single-cell level, have been proposed to play an important role in early stage embryonic stem cell differentiation (*19–21*) and epigenetic mechanisms are increasingly believed to play an important role the development, progression and emergence of drug resistance in cancer (*22, 23*). Furthermore, it may have implications for important biotechnologies, such as reprogramming of human somatic cells to generate induced Pluripotent Stem Cells (iPSCs) (*24*).

## Acknowledgments

The authors thank Paul M. Chaikin (New York University) for helpful discussions.

## Author contributions

G.W., P.N., J.Baudry, L.B., E.B., A.D.G. and J.Bibette designed the study; G.W. performed the experiments; G.W. and P.N. analyzed the data; G.W., P.N., J.Baudry, E.B., A.D.G. and J.Bibette interpreted the data and wrote the paper.

## Data and materials availability

Supplementary information is available for this paper. Correspondence and requests for materials should be addressed to A.D.G. or J.B.

## Materials and Methods

All chemicals were purchased from Sigma-Aldrich unless otherwise mentioned.

### Strains and culture conditions

The *Saccharomyces cerevisiae* YPH499 strain carries a deletion of the endogenous chromosomal *HIS3* gene and a plasmid containing the divergent *GAL1/GAL10* promoters. There is a *HIS3* gene under sole regulation of *GAL1* promoter, and a *GFP* reporter gene under control of the *GAL10* promoter (*4*). Cells were grown from colonies on plates in 25 mL of culture medium comprising: 1.7 g/L of yeast nitrogen base without amino acids and ammonium sulfate, 5 g/L ammonium sulfate, 1.4 g/L amino acid dropout powder (without Trp, His, Leu, Ura), 0.004 g/L L-tryptophan, and 0.002 g/L uracil, 20 g/L of galactose or glucose; and incubated at 30°C, shaking at 200 r.p.m. in 100 mL Erlenmeyer flasks. Cultures were diluted every 12h to maintain OD<1.0 (never transferring less than 3.10^6^ cells).

Rewired yeast cells, were first grown in a batch culture with galactose medium lacking histidine. The optical density (OD600nm) of the culture was monitored and it was maintained in the exponential phase of growth by serial dilutions for 2 days. The medium was then switched from galactose to glucose. As previously reported (*4, 9*), naïve cells (i.e. rewired cells that had never been grown in glucose before) were able to grow for about 20 hours in glucose media lacking histidine with a doubling time, *t*_D_, of 1.9 h (phase I). Following this phase, growth slowed considerably (*t*_D_ 17.1 h), leaving the OD_600nm_ approximately constant (phase II, the “latent phase”). At the end of this phase, after durations that varied across repeated experiments from (1 day to 6 days), the population adapted and started growing and proliferating (phase III, the “adapted phase”). The doubling time of the adapted population reduced from 3.1 h 1 day after entering phase III, to 2.1 h after 8 days in phase III, reflecting the fact that the population continues to adapt after resuming growth (*5*). Three independent batch cultures were analyzed and the growth curve for one of these experiments is shown in Fig. 1.

### Microfluidic device fabrication

Microfluidic devices were obtained using conventional soft lithography methods (*25*), as described (*26*). Molds were prepared using SU8-2015 or SU8-2075 photoresist (MicroChem Corp.) and used to pattern 20 and 75 μm-deep channels onto silicon wafers (Siltronix). The channels of the devices were passivated with Aquapel in HFE7100 (3M) and subsequently flushed with compressed nitrogen gas.

### Formation and imaging of droplet arrays

Single-cell metabolic and growth dynamics of yeast cells from batch cultures were measured as in Boitard et al. (*11*). Yeast cells from batch culture were individually compartmentalized in monodisperse 30 pL volume (38 μm diameter) aqueous droplets containing fresh glucose medium lacking histidine by hydrodynamic flow-focusing (*27*) with a fluorinated oil phase containing fluorosurfactant (*28*). For each experiment, about 200,000 droplets were incubated at 30°C in a 2D array and images were taken every 30 minutes over three days. Briefly, a flow-focusing device (*27*) was used for droplet generation and flow rates were controlled using standard-pressure infuse/withdraw PHD 22/2000 syringe pumps (Harvard Apparatus Inc., Holliston, MA). Syringes (Hamilton) connected to the microfluidic device using 0.6 x 25 mm Neolus needles (Terumo Corporation) and PTFE tubing with an internal diameter of 0.56 mm and an external diameter of 1.07 mm (Fisher Bioblock Scientific). The aqueous phase, comprising cells suspended in culture medium, was injected at 300 μL/h and dispersed in a continuous phase consisting of HFE-7500 fluorinated oil (3M) containing 2% (w/w) EA surfactant (Raindance Technologies), a PFPE-PEG-PFPE amphiphilic tri-block copolymer (28), injected at 150 μL/h, forming 30 pL volume (38 μm diameter) droplets at 2,800 droplets s^-1^. Droplets were produced in a compact manner and were directly incubated in a glass chamber of 3.5 x 1.5 cm to form a compact 2D droplet array (11). Compaction of the emulsion prevents droplet movement to enable their tracking over long time scales. The chamber was maintained at 30°C on a Nikon T300 inverted microscope with a Thorlabs MAX202 XY stage. Images of the droplet array were taken every 30 min using a Hamamatsu Orca-ER camera. Custom-made Labview software was used to automate image acquisition and microscope control.

### Data processing

For a schematic representation see Fig. S5.

### Droplet tracking and normalization

Droplets were detected with an in-house Matlab segmentation routine and tracked over time by a nearest neighbour criterion, knowing that displacements from image to image are much smaller than the characteristic droplet diameter. Images in which large displacements occured were easily detected as they displayed strongly discontinuous volume traces and were discarded. In order to minimize volume fluctuations caused by defocusing in time, volumes were normalized by the time course of the average of more than 10 empty droplets for each image. Images for which normalization failed were discarded (Fig. S5). The first and second derivatives of the normalized volume over time were computed over 5 time point sliding windows and provide the metabolic rate and metabolic acceleration and were used to automatically determine the maximum metabolic rate, time of metabolic acceleration and time of metabolic arrest (Fig. S3).

### Significantly shrinking droplets

A droplet was considered to be significantly shrinking (i.e. the compartmentalized cell showed detectable metabolism) if its final normalized volume *V*_end_ was smaller than *V*_s_ =1-3σ, where σ is the coefficient of variation of empty droplet final volumes taken over all experiments. We measured *V*_s_ = 0.95. Shrinking but empty droplets that passed this filter were removed by visual inspection of the droplet content.

### Poisson parameter and initial fraction of dead cells

The Poisson parameter *λ* (the mean number of cells per droplet) was determined from the frequency of occupied droplets on a sample of 10 initial images (~1500 droplets) per experiment. The number of droplets containing cells arrested from the start was estimated by calculating the difference between *λ* and the number of significantly shrinking droplets (Fig. S1).

### Metabolic arrest

Metabolically arrested cells were defined as cells: (i) which do not consume all the nutrients available in the droplets within the timeframe of the experiment, as defined by *V*_end_ > *V*_e_, where *V*_end_ is the final droplet volume and *V*_e_ = 0.55 is the maximal normalized volume reached by cells in the control experiment (with histidine), and (ii) for which the volume trace significantly flattens during the time course of the experiment. The latter was determined as the existence of an inflection in the volume curve leading to reach a rate *R* < *R*_max_/10, where *R*_max_ is the maximum rate observed for this cell. The metabolic arrest point is defined as the time at which *R* reaches *R*_max_/10 (Fig. S3). In the presence of histidine, an abrupt decrease in metabolic rate was reached when glucose in the droplet was exhausted by the cells (*11*), corresponding to a mean final droplet volume of 0.49±0.002 of the initial value, and always <0.55 of the initial value (Fig. 2F, red). In contrast, in the adaptation experiment in the absence of histidine (Fig. 2F, blue), 54% of cells had a final volume >0.55 of the initial value, indicating that they had not consumed all the glucose in the droplet. These cells fall into two categories. In the first category are cells (49%) that arrested their metabolism during the experiment (Fig. 2G), as they were initially metabolically active and showed an abrupt metabolic deceleration and a stable volume, indicating that the cells were not metabolically active from that point in time (*T*_arr_). A second category of cells (5%) neither arrested, nor recovered, but had a metabolism slow enough that glucose was not exhausted at the end of the experiment.

### Metabolic recovery

Metabolic recoveries in the adaptation experiment were determined as those cells whose metabolic rate increased to levels comparable to the control with histidine, as computed by the difference (Δ*R*) between the maximum rate (*R*_max_) and the initial rate (*R*_0_) (2 hours after compartmentalization) obtained from the volume time course of each droplet (Fig. 2C). For this we determined the lower and upper bounds of the crossing values for the histograms of *ΔR* (Fig. 2H) for different binning intervals in the presence and absence of histidine (Fig. S4). The threshold value *ΔR*\* = 0.013 h^-1^, which indicates the cross-over between the two distributions, was obtained when these bounds converged. Cells with *ΔR* > *ΔR*\* were classed as metabolic recoveries. In the control, with histidine, the metabolic rate of cells steadily increased due to cell growth and division before glucose exhaustion (Fig. 2B) (*11*), with a mean rate increase of 3.6±0.7 10^-2^ h^-1^ (Fig. 2H, red). In the adaptation experiment, 4% of cells accelerated their metabolic rates (N=120, Fig. 2H, blue) to values similar to those observed in the control with histidine (>1.3 10^-2^ h^-1^).

### Computation of instantaneous rates of arrest and metabolic recovery

Three distinct cell fates were categorized from the metabolic curves: metabolic arrest, metabolic recovery and steady metabolism. Calling *N* the total number of cells observed during the experiment, *N_a_* and *N_r_* the respective cumulative number of arrested and recovering cells, the instantaneous rate of arrest and recovery were respectively computed as *dN_a_*/*dt*/(*N* – *N_a_* – *N_r_*) and *dN_r_*/*dt*/(*N* – *N_a_* – *N_r_*).

### Effective coefficients of determinations

The correlations of Fig. 4C,H display regions of experimentally inaccessible parameter values, due to the finite amount of resource available in each droplets and the noise in droplet volume measurements. The coefficient of determination *R*_a_^2^ computed directly from the data thus comprises a contribution of these boundary constraints. This contribution *R*_b_^2^ to the total covariance was estimated using a Monte Carlo approach preserving the marginal distributions: 10^5^ random permutations of the measured values were generated under the boundary constraints, taking *R*_b_^2^ as the mean coefficient of determination of the last 10^4^ realizations. Assuming independence between the experimental constraints and cell fates in the absence of resource exhaustion, the additive contribution of these phenomena to the covariance leads to effective coefficients of determinations *R*^2^= *R*_b_^2^ – *R*_a_^2^.

**Fig. S1.**
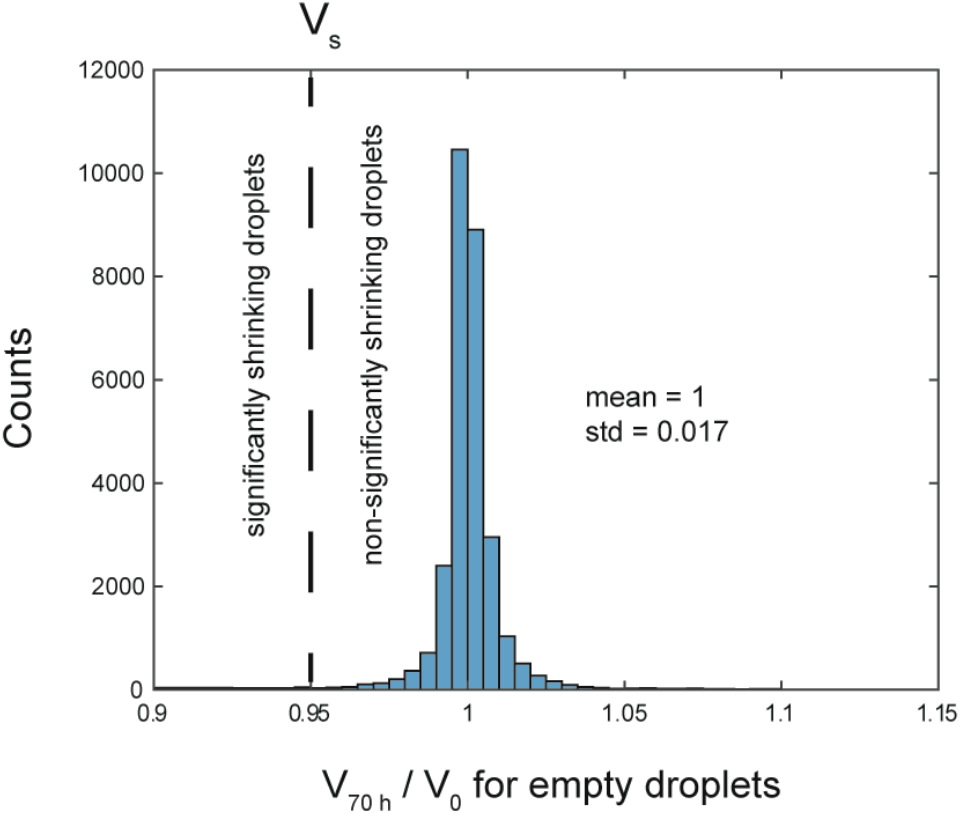
Threshold for significant shrinking. The threshold *V*_s_ = 0.95 for maximal final volume indicating a significant volume decrease is defined as 3 standard deviation away from the mean of volume distribution of empty droplets.

**Fig. S2.**
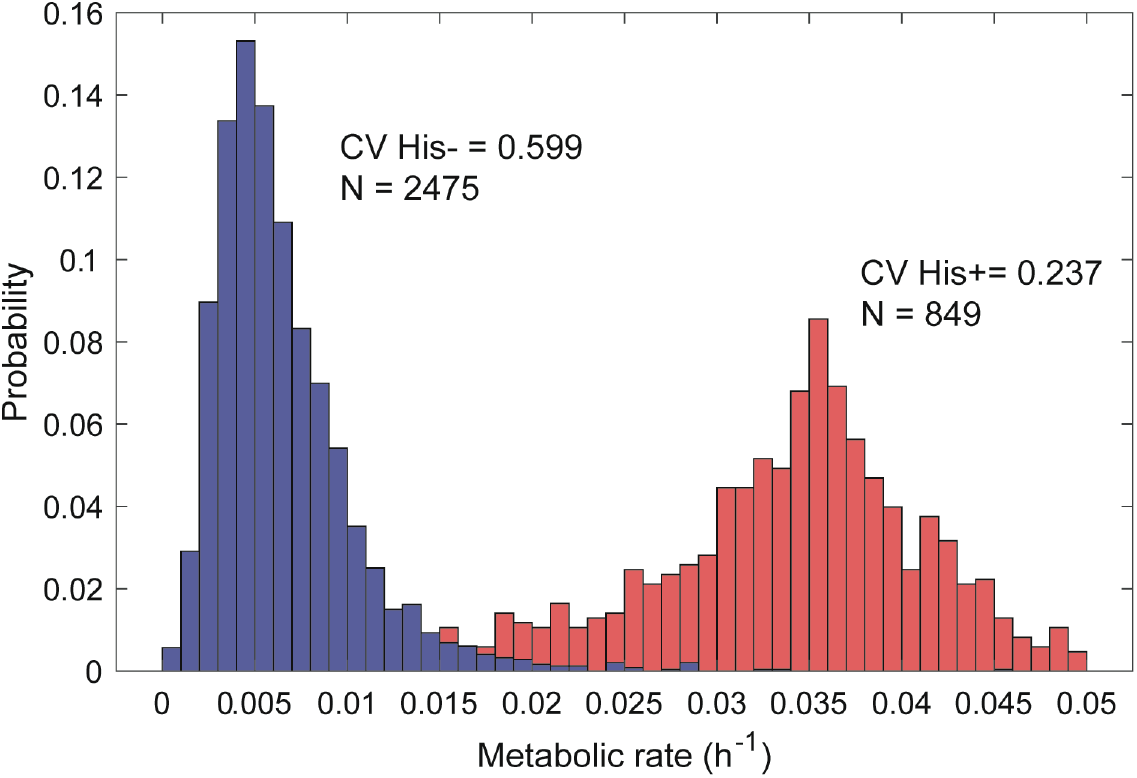
Distribution of metabolic rates at 10 hours after encapsulation of yeast in droplets. Red: in the absence of metabolic challenge (with histidine). Blue: in response to a metabolic challenge (no histidine).

**Fig. S3.**
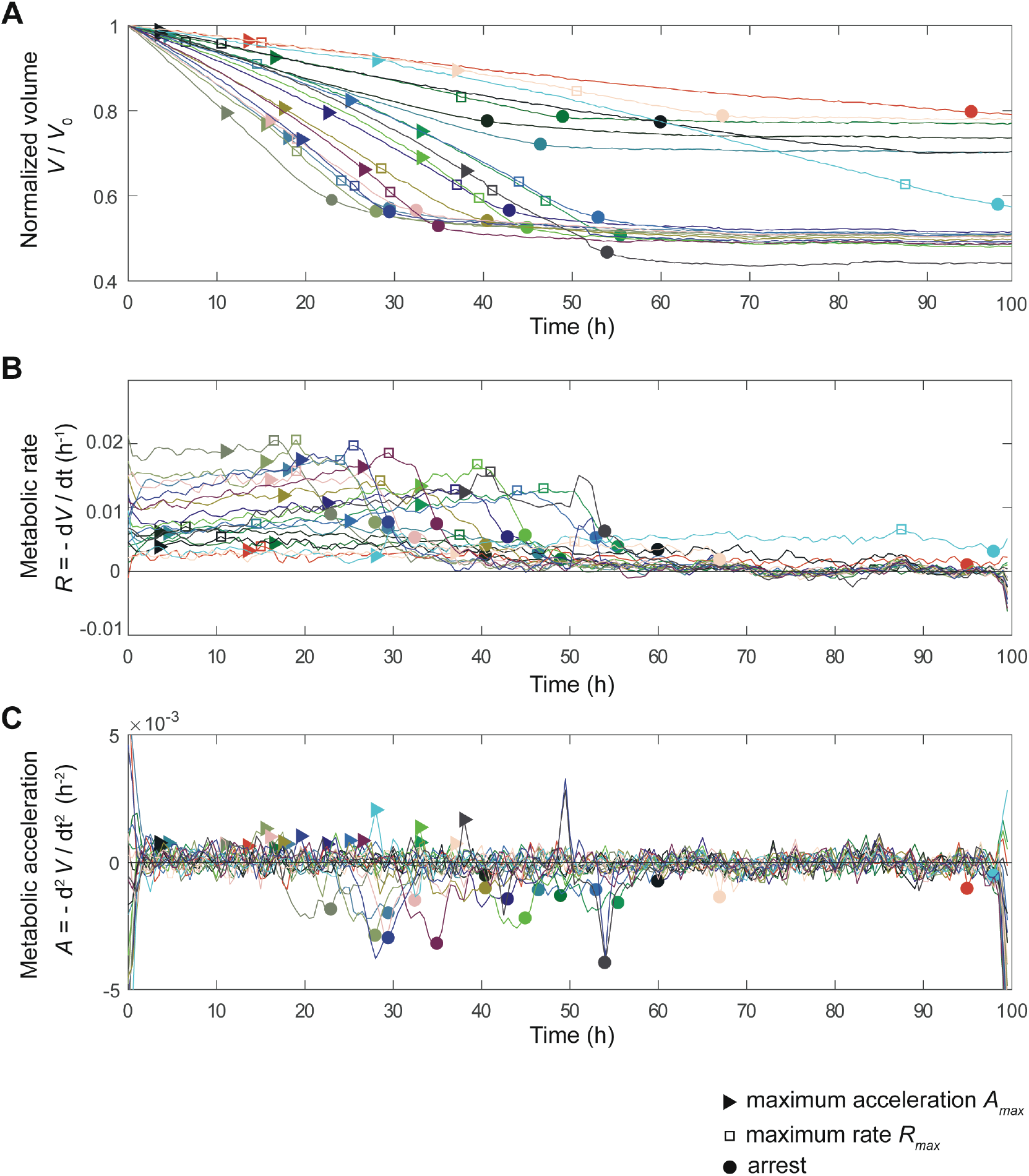
Automated analysis of metabolic kinetics from the time-lapse measurement of droplet volumes. A) Example of 20 droplet volumes *V* over time normalized by their initial volume, V0. The point of metabolic acceleration (*A_max_*) is shown by triangles, of maximum rate (*R_max_*) by squares and metabolic arrest (Tarr) by circles (see below). B) Metabolic rate *R* computed as minus the time derivative of the normalized volume. *R_max_* (square) is determined as the maximum of each curve. Metabolic arrest is determined as the point at which *R* reaches 1/10 of *R_max_*, and generally corresponds to the minimum of the acceleration curves in panel c, although the latter yields a less reliable estimate. C) Metabolic acceleration *A* computed as the derivative of the metabolic rate. The time of acceleration (triangle) is determined from the maximum *A_max_* of this curve. The time of recovery (Trec) is the time at which *A_max_* is reached within the population of cells that significantly increase their metabolism (ΔR > ΔR*) (see Fig. 2H and Fig. S4).

**Fig. S4.**
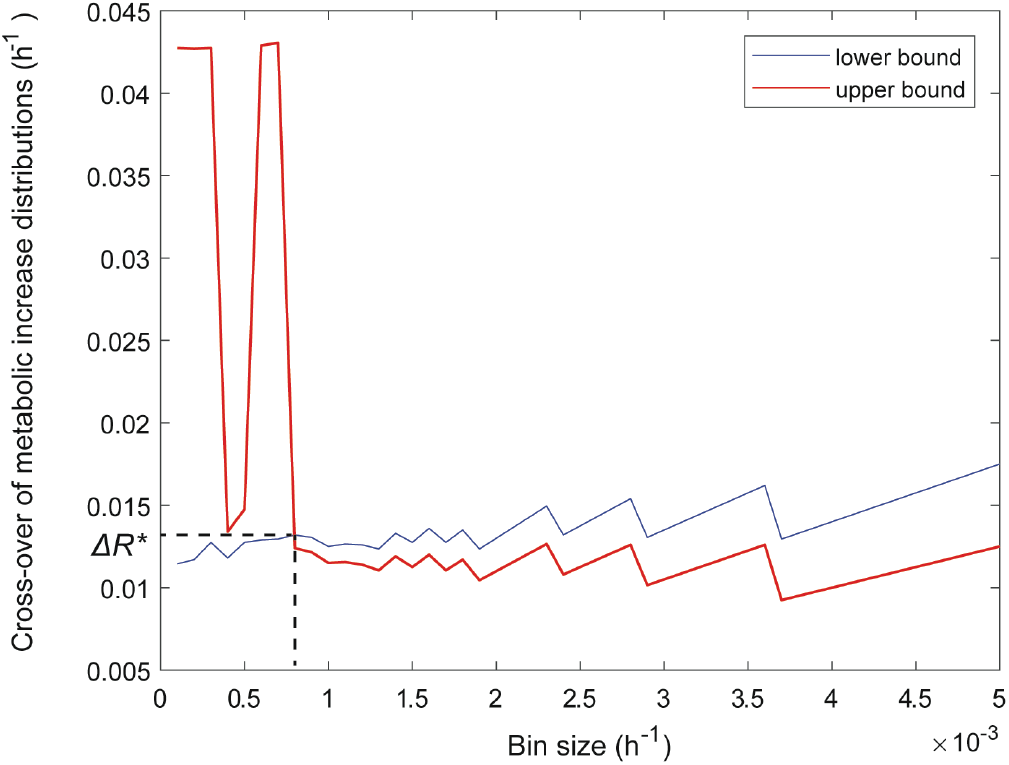
Determination of the metabolic increase threshold *ΔR*\* for significant metabolic recovery. The metabolic increase *ΔR* is the difference between the initial metabolic rate (taken at 2 hours) and the maximum measured metabolic rate *R_max_* (see Fig. S3). For each bin size, we determined the lower and upper bound for the cross-over between the metabolic increase distributions (see Fig. 2H), in the presence or absence of metabolic challenge. These bounds converge at a certain bin value which optimally defines the cross-over *ΔR*\* between these two distributions.

**Fig. S5.**
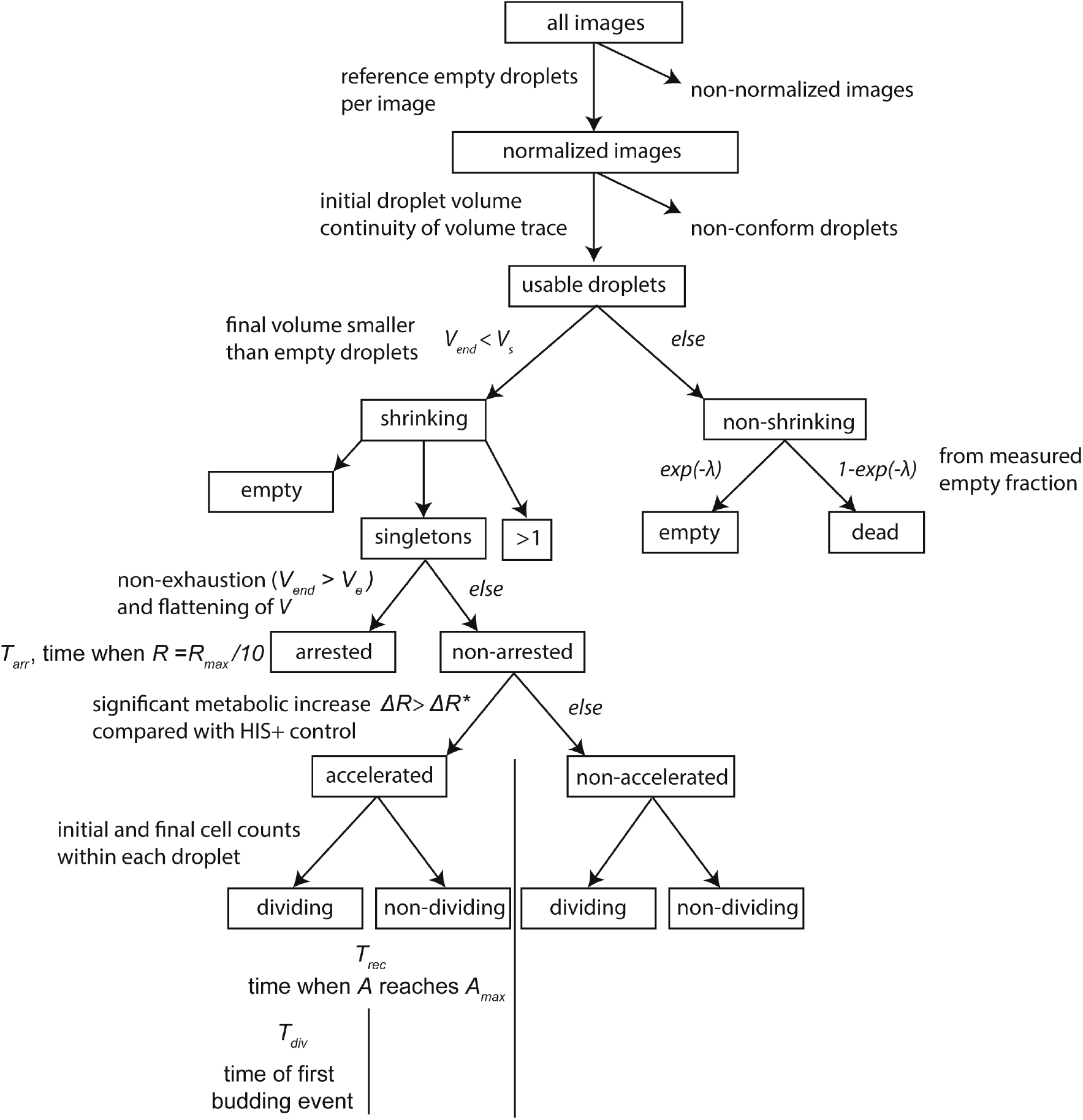
Classification tree of metabolic responses. For the details of the treatment, refer to the Materials and Methods. *V_end_* is the final droplet volume; *V*_s_ =1-3σ, where σ is the coefficient of variation of empty droplet final volumes taken over all experiments; *V_e_* is the maximal normalized volume reached by cells in the control experiment (with histidine); *λ* is the Poisson parameter corresponding to the average number of cells encapsulated per droplet; *R* is the metabolic rate; *R_max_* is the maximum metabolic rate reach for each cell; *T_arr_* is the time at which *R* reaches *R_max_/10* in arresting cells; *ΔR* is the metabolic rate increase computed as the difference between *R_max_* and the initial rate *R*_0_ (at 2 hours); *ΔR*\* is the cross-over value between low and high metabolic rate increases (Fig. S4); *A* is the metabolic acceleration, which is the time derivate of *R*; *A_max_* is the maximum metabolic acceleration (Fig. S3); *Trec* is the time at which *A_max_* is reached within the population of cells that significantly increase their metabolism (*ΔR* > *ΔR*\*); *T_div_* is the time at which the first bud appears in droplets where the final number of cells is larger than 2.

**Fig. S6.**
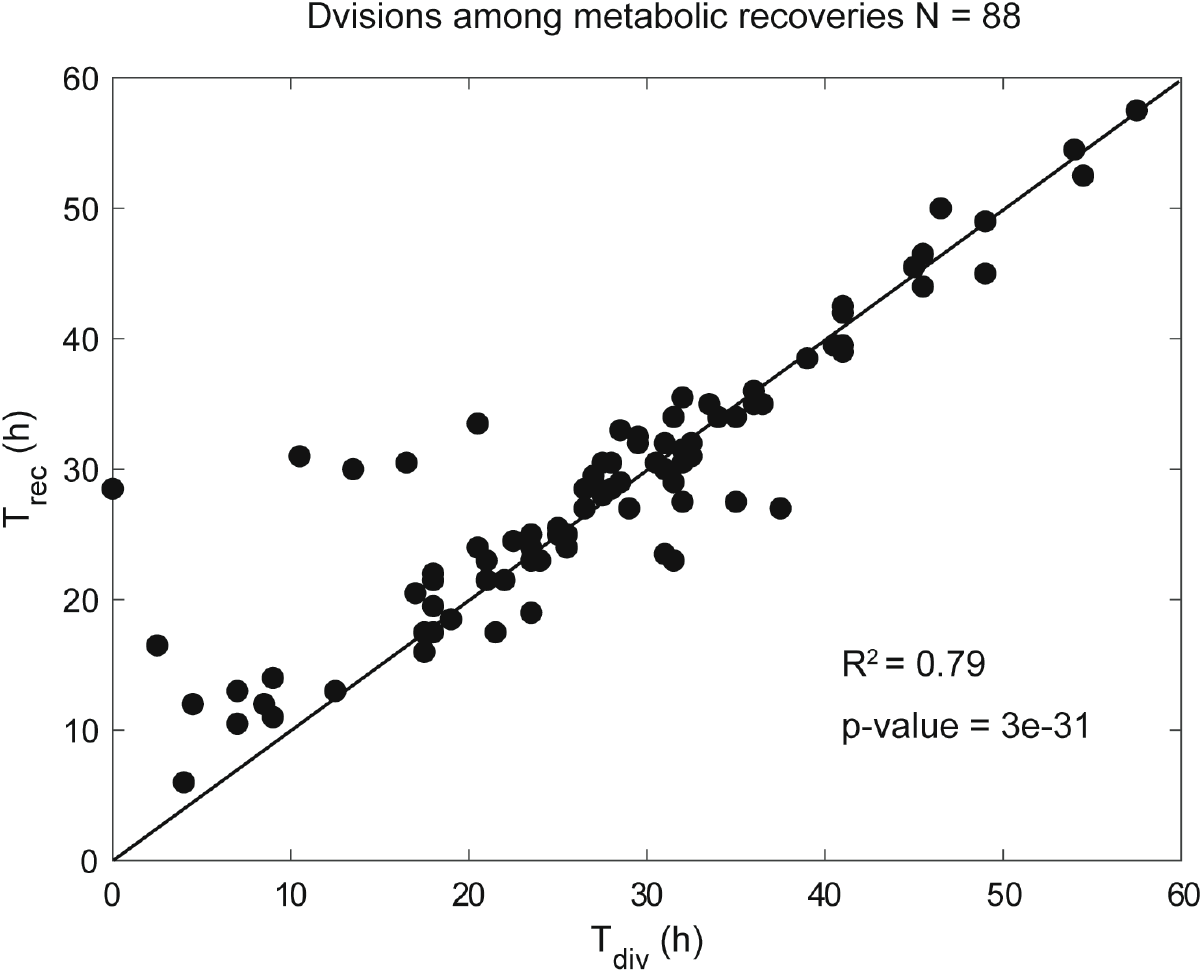
Correlation between the times of division (*T*_div_) and metabolic recovery (*T*_rec_). Each dot corresponds to the analysis of a single volume trace measured in the absence of histidine. The diagonal corresponds to *T*_div_ = *T*_rec_.

**Fig. S7.**
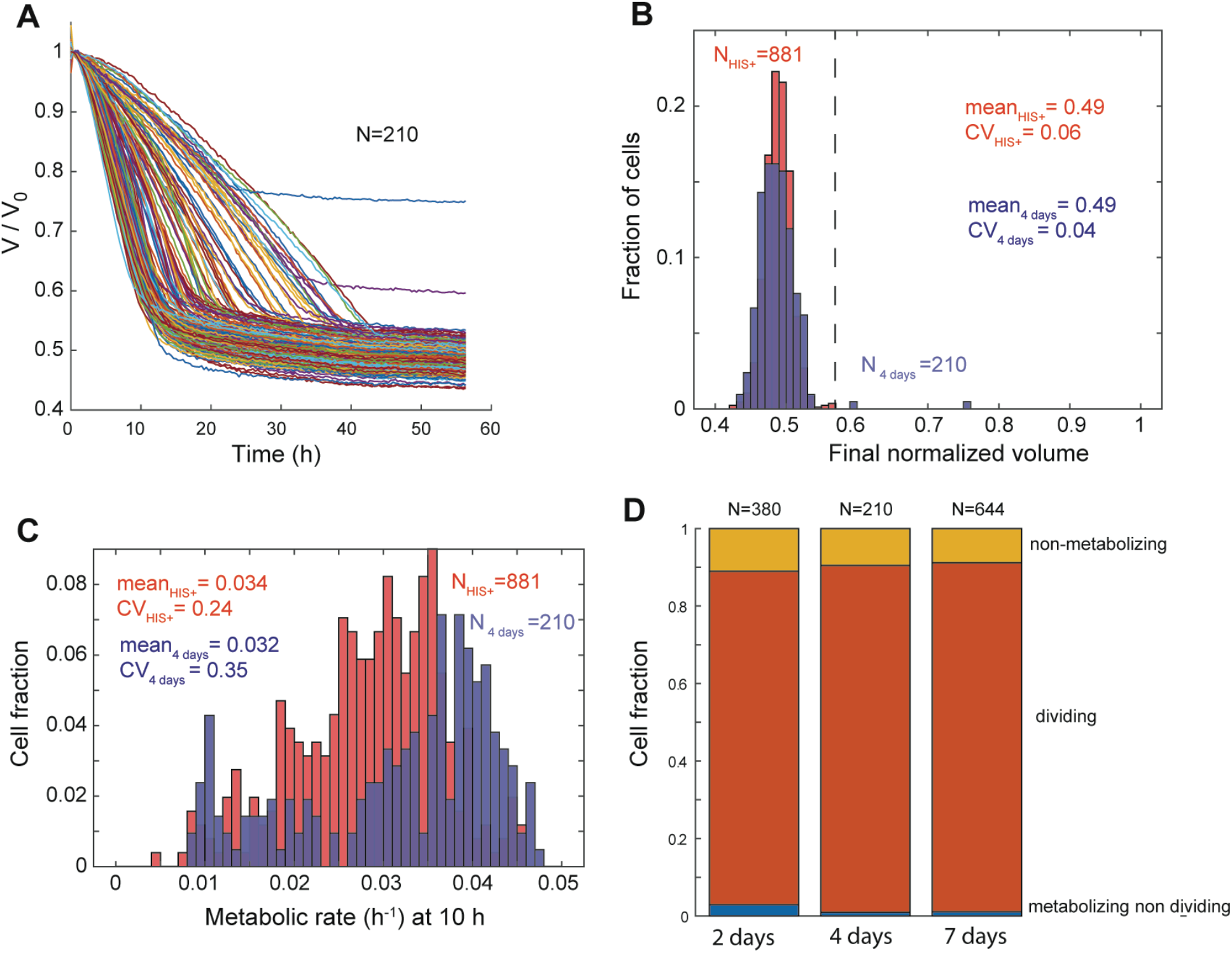
Adapted population. A) Volume traces of droplets containing single cells taken from the culture 4 days after the challenge. Droplet volume *V* is normalized by the initial volume *V_0_*. Lines are randomly color-coded. B) Distribution of final volumes (at 70 hours) 4 days after the challenge (blue) and for the control with histidine (red). The vertical dashed line represents *V_e_*, the maximum normalized volume reached by cells in the histidine positive control. C) Distribution of metabolic rates at 10 hours, 4 days after stress (blue) and for the control with histidine (red). The variance in metabolic rate of the dividing cells is similar in the samples taken 2, 4 and 7 days after entering the adapted phase (CV = 0.35) and close to that observed for control cells switched to glucose plus histidine (CV = 0.24). D) Fraction of each cell type 2, 4 and 7 days after the challenge. The proportions of the different classes of cells are very similar at the three time points: non-metabolizing cells (1.6±1%), non-dividing metabolizing cells (10±1%) and dividing cells (89±2%). Consistent with adaptation observed at the population level, the fraction of dividing cells is >10-fold higher than at the beginning of Phase II (5).

**Table S1. Parameters measured and computed for each individual experiment after stress, after adaptation, for the control with histidine, and over groups of experiments**. In all cases, the symbol “N” refers to a number of droplets.

**Movie S1. Droplets were incubated at 30°C in a sealed glass chamber imaged over 70 h**. Images were taken every 30 min, as indicated by the time stamp. Individual cells were compartmentalized in aqueous droplets dispersed in an oil phase and parked in a 2D array in a closed glass chamber. About 6% of the droplets contain a single cell. The consumption of nutrients (glucose) in a cell-containing droplet creates an osmotic imbalance, resulting in an osmotically driven water flux, which induces the shrinkage of this droplet and the swelling of neighboring cell-free droplets The change in volume of the droplet containing the cell is extracted by image analysis to measure cell metabolism as a function of time, while imaging enables measurement of cell division.

