## Supplementary material for "Metabolic Cost of Rapid Adaptation of Single Yeast Cells": Table S1

| | Right after stress | Right after stress | Right after stress | 2 days after adaptation | 4 days after adaptation | 7 days after adaptation | Control with histidine | Consolidation stress | All |
|---|---|---|---|---|---|---|---|---|---|
| **N images all** | 300 | 300 | 300 | 96 | 96 | 96 | 96 | 900 | 1284 |
| **N droplets all** | 109522 | 144619 | 75380 | 39001 | 43865 | 23054 | 24814 | 329 521 | 460255 |
| **fraction occupied droplets** | 0,076 | 0,041 | 0,051 | 0,038 | 0,045 | 0,080 | 0,130 | 0,05 | NA |
| **Poisson parameter λ** | 0,079 | 0,042 | 0,052 | 0,039 | 0,046 | 0,083 | 0,139 | 0,06 | NA |
| **N images normalized** | 199 | 174 | 224 | 96 | 62 | 96 | 88 | 597,00 | 939 |
| **N droplets normalized** | 28013 | 18846 | 31413 | 35754 | 11765 | 18922 | 7917 | 78 272,00 | 152630 |
| **N droplets usable** | 26240 | 17865 | 30829 | 35750 | 11691 | 18733 | 7845 | 74 934,00 | 148953 |
| **inferred N empty** | 24246 | 17133 | 29257 | 34378 | 11164 | 17242 | 6827 | 70 635,02 | 140246 |
| **std( $V_{end}$ / $V_0$) for empty droplets** | 0,0126 | 0,0091 | 0,0174 | 0,0069 | 0,0081 | 0,0175 | 0,0184 | NA | NA |
| **$V_s$ / $V_0$** | 0,95 | 0,95 | 0,95 | 0,95 | 0,95 | 0,95 | 0,95 | 0,95 | 0,95 |
| **N shrinking** | 1441 | 681 | 1659 | 1127 | 513 | 1483 | 929 | 3 781,00 | 7833 |
| **N non-shrinking** | 24799 | 17184 | 29170 | 34623 | 11178 | 17250 | 6916 | 71 153,00 | 141120 |
| **N containing dead cells** | 553 | 51 | 0 | 245 | 14 | 8 | 89 | 518 | 960 |
| **N containing cells** | 1994 | 732 | 1659 | 1372 | 527 | 1491 | 1018 | 4 299 | 8793 |
| **Fraction of dead cells** | 27,7% | 7,0% | 0,0% | 17,8% | 2,7% | 0,5% | 8,8% | 0,12 | 10,9% |
| **$V_e$ / $V_0$** | 0,55 | 0,55 | 0,55 | 0,55 | 0,55 | 0,55 | 0,55 | 0,55 | 0,55 |
| **inferred total cell population** | 1340 | 559 | 987 | 463 | 216 | 647 | 931 | 2814 | 5117 |
| **N discarded** | 472 | 160 | 671 | 747 | 303 | 839 | 80 | 1303 | 3272 |
| **N all checked shrinking** | 968 | 520 | 987 | 380 | 210 | 644 | 849 | 2475 | 4558 |
| **N non exhausted** | 391 | 423 | 699 | 10 | 2 | 9 | 3 | 1513 | |
| **N arrest** | 385 | 379 | 604 | <11 | 2 | 7 | 2 | 1368 | 1379 |
| **N accelerated** | 38 | 25 | 57 | NA | NA | NA | NA | 120 | 120 |
| **N non accelerated** | 545 | 116 | 326 | NA | NA | NA | NA | 987 | 987 |
| **N accelerated dividers** | 23 | 22 | 43 | NA | NA | NA | NA | 88 | 88 |
| **N non accelerated dividers** | 28 | 16 | 41 | NA | NA | NA | NA | 85 | 85 |
| **N dividing** | 51 | 38 | 84 | 327 | 188 | 580 | 847 | 173 | 2115 |
| **mean metabolic rate at 10 h ($h^{-1}$)** | 8,34E-03 | 4,32E-03 | 5,81E-03 | 2,76E-02 | 3,24E-02 | 3,85E-01 | 0,0335 | 6,49E-03 | NA |
| **CV metabolic rate at 10 h** | 52% | 53% | 55% | 30% | 35% | 39% | 24% | 60% | NA |
| **mean rate increase ΔR ($h^{-1}$)** | 6,14E-03 | 7,92E-03 | 8,02E-03 | NA | NA | NA | 0,037 | 7,02E-03 | NA |
| **std ΔR (h-1)** | 4,72E-03 | 7,77E-03 | 6,56E-03 | NA | NA | NA | 0,0069 | 5,92E-03 | NA |
| **maximum ΔR ($h^{-1}$)** | 4,27E-02 | 3,74E-02 | 3,49E-02 | NA | NA | NA | 0,0628 | 4,27E-02 | NA |
| **$T_{div}$ - $T_{acc}$ (h)** | -1,50E+00 | -2,05E-01 | -1,86E+00 | NA | NA | NA | NA | -4,51E+00 | NA |
| **std($T_{div}$ - $T_{acc}$) (h)** | 6,95E+00 | 2,72E+00 | 5,70E+00 | NA | NA | NA | NA | 1,33E+01 | NA |
| **$R^2$ correlation $T_{div}$ $T_{acc}$** | 7,36E-01 | 9,47E-01 | 7,08E-01 | NA | NA | NA | NA | 3,60E-01 | NA |
| **p-value correlation** | 8,35E-08 | 3,02E-14 | 2,94E-12 | NA | NA | NA | NA | 6,27E-18 | NA |
| **divinding fraction among metabolic recoveries** | 6,05E-01 | 8,80E-01 | 7,54E-01 | NA | NA | NA | NA | 7,33E-01 | NA |
| **quadratic rate of arrest ($h^{-2}$)** | 7,95E-05 | 1,11E-04 | 1,19E-04 | NA | NA | NA | NA | 1,02E-04 | NA |
| **95% confidence** | 7.79e-05 8.10e-05 | 1.08e-04 1.13e-04 | 1.18e-04 1.20e-04 | NA | NA | NA | NA | 1.01e-04 1.02e-04 | NA |
| **quadratic rate of metabolic recovery ($h^{-2}$)** | 3,09E-05 | 2,09E-05 | 2,92E-05 | NA | NA | NA | NA | 2,75E-05 | NA |
| **95% confidence** | 2.95e-05 3.23e-05 | 1.71e-05 2.48e-05 | 2.78e-05 3.06e-05 | NA | NA | NA | NA | 2.66e-05 2.83e-05 | NA |
| **quadratic rate of division recovery ($h^{-2}$)** | 1,85E-05 | 1,60E-05 | 2,73E-05 | NA | NA | NA | NA | 2,23E-05 | NA |
| **95% confidence** | 1.68e-05 2.02e-05 | 1.30e-05 1.90e-05 | 2.66e-05 2.80e-05 | NA | NA | NA | NA | 2.19e-05 2.28e-05 | NA |
| **$R^2$ correlation $T_{rec}$ vs $1/R_{ini}$** | 0,75 | 0,07 | 0,26 | NA | NA | NA | NA | 3,31E-01 | NA |
| **p-value correlation** | 4,17E-13 | 1,94E-01 | 5,19E-05 | NA | NA | NA | NA | 2,37E-12 | NA |
| **variance explained by inequality constraints** | 2,21E-02 | 1,91E-06 | 1,41E-03 | NA | NA | NA | NA | 1,04E-02 | NA |
| **mean $V_{rec}$ / $V_0$** | 0,72 | 0,78 | 0,74 | NA | NA | NA | NA | 0,74 | NA |
| **$R^2$ correlation $T_{arrest}$ vs $1/R_{ini}$** | 0,14 | 0,12 | 0,07 | NA | NA | NA | NA | 1,17E-01 | NA |
| **p-value correlation** | 4,89E-22 | 1,14E-10 | 5,68E-13 | NA | NA | NA | NA | 3,19E-48 | NA |
| **variance explained by inequality constraints** | 0,11 | 0,00 | 0,01 | NA | NA | NA | NA | 4,14E-02 | NA |
